# Environmental Selection Shapes Resistance, Metabolic, and Adaptive Capabilities in Exiguobacterium

**DOI:** 10.64898/2026.02.03.703531

**Authors:** Monserrat Manzo-Ruíz, Patricia Espinosa Cueto, Janeth Valdés-Hernández, Jenifer Sánchez-López, Luis Daniel Ríos-Becerra, Alba Romero Rodríguez

## Abstract

The genus *Exiguobacterium* comprises Gram-positive, non-spore-forming, facultative anaerobic bacteria known for their remarkable adaptability to extreme environments, including soils, hot springs, glaciers, and the gastrointestinal tracts of certain organisms. Despite their unique adaptations for surviving in extreme environments, their pathogenicity is well documented. Here, we analyzed the phenotypical traits of two Mexican strains of *Exiguobacterium*—JVH47, isolated from contaminated urban sediments in Mexico City, and P4526, from the less human-impacted Cuatro Ciénegas Basin. Furthermore, strains were related via comparative genomics using publicly available genomes.

Phenotypic characterization demonstrated that both strains thrive across a wide range of temperatures (20–50 °C), pH (7–11), and salinity (up to 7% NaCl). Although sensitive to erythromycin, the JVH47 strain exhibited higher erythromycin resistance and harbored antibiotic resistance genes. This study underscores the ecological versatility of *Exiguobacterium* and its potential role as a reservoir for antibiotic resistance genes. While rarely associated with human infections, its ability to survive in extreme conditions and form biofilms raises concerns for immunocompromised individuals. These findings highlight the need for careful consideration of *Exiguobacterium* in biotechnological applications and its implications under the One Health framework.

## Introduction

The *Exiguobacterium* genus consists of Gram-positive, non-spore-forming, and facultative anaerobic bacteria. This genus comprises halophilic, psychrotrophic, and moderate thermophilic species isolated from many environments, including soils, rhizosphere, hot springs, glaciers, and even permafrost [1]. Furthermore, *Exiguobacterium* has been isolated from the gastrointestinal tract of fish, crustaceans, and mealworms [2–4]. This enormous inhabitability may rely on their wide range of metabolic and stress resistance capabilities, including genes encoding multiple carbohydrate hydrolases, comprising amylases and chitinases [1, 4]. The metabolic capability of the *Exiguobacterium* genus has been explored for industrial and biotechnological applications, bioremediation, and agriculture [2, 5]. Recently, strains of the genus *Exiguobacterium* have been introduced with plant growth-stimulating attributes for growth promotion in several crops [5].

Furthermore, *Exiguobacterium* members have been proposed as potential probiotics to enhance production in crustaceans and fish [6–8]. Nevertheless, under the One Health approach, significant considerations should be made when using some strains as probiotics for human-animal consumption or when delivering strains in the environment. It is well known that many saprophytic soil bacteria are drug-resistant and that those genes could be mobilized to human pathogens. Antibiotic resistance profiles or the risk for mobilization in this genus are largely unknown. The *Exiguobacterium* sp. strain S3-2, isolated from sediment at a fish farm, exhibited a multidrug-resistant phenotype. Five resistance genes showed significant similarity (83%-100%) to sequences found in pathogens [17]. The authors suggest that, for that strain, acquisition of the antibiotic resistance gene was likely associated with antibiotics used in fish farms [17].

Furthermore, although rare, *Exiguobacterium* strains have been associated with bacteremia, skin infections, and even a fatal case of community-acquired pneumonia in humans [9–12]. Even though most of the cases of infections associated with the *Exiguobacterium* genus have been diagnosed in immunocompromised patients or associated with specific risk factors, such as drug abuse [12]. However, the increasing number of environmental saprophytic bacterial species that can cause infections in immunocompromised hosts and individuals of advanced age is a serious concern.

In this paper, we aim to compare two Mexican strains: one isolated from contaminated soil in a megapolis and the other from ancient soil, to examine their adaptive traits. Furthermore, comparative genomics was used to address antibiotic resistance and potential virulence.

## Materials and Methods

### Bacterial strains and culture conditions

*Exiguobacterium sp.* JVH47 was isolated from sediments of an urban flood in Mexico City. *Exiguobacterium sp*. P4526 was isolated from Cuatro Ciénegas Basin [13]; and it was kindly donated by Dra. Valeria Souza. Strains were routinely grown in Brain Heart Infusion (BHI), BD™. For assays, the strains were grown on Luria-Bertani (LB) (Tryptone 10.0 gL-1, Yeast extract 5.0 gL-1, NaCl 10.0 gL-1, pH 8.5). For cytotoxic assays, bacteria were grown in Dulbecco′s Modified Eagle Medium (DMEM), Gibco™, for 48 and 72 h.

### Identification of bacterial species

For identification, DNA was isolated conventionally by the phenol-chloroform method, and then the 16S rRNA gene sequence was amplified by PCR using Primers 16S rRNA For (5’-AGA GTT TGA TCC TGG CTC AG-3’) and 16S rRNA Rev (5’-ACG GCT ACC TTG TTA CGA CTT-3’). PCR reaction mixture containing Taq polymerase (ABClonal®) was set under the following cycling conditions: 94 °C for 2min, (94 °C for 30 s, 60 °C for 30 s, 74 °C for 1 min) X25 cycles, 74 °C for 5 min PCR amplified products were purified using NZYGelpure (NZYTech) and sent for sequencing at the Laboratory of biodiversity genomic sequencing, UNAM (Mexico). Sequence similarity was verified using the Basic Local Alignment Search Tool (BLAST) v2.17.0 at NCBI (https://blast.ncbi.nlm.nih.gov/Blast.cgi). Sequences of both isolates, *Exiguobacterium* sp. strain JVH47 and P4526, have been submitted to NCBI

### Determination of temperature, pH, and salinity limits of strain survival

To assess the range of adaptation to pH, temperature, and salinity, strains were monitored for growth under the following conditions: for temperature-based assessments at 4, 20, 29, 37, 40, and 50 °C on LB broth pH 8.5; for pH-based assessments, at 37 °C in liquid LB medium in 1-unit steps from pH 4.0 to pH 11.0; and for salinity assessments, from 0 to 20% w/v, NaCl at 37 °C. Growth was assessed by measuring optical density at 600 nm (OD_600_) at 24, 48, and 72 h.

### Cell survival in the presence of erythromycin

*Exiguobacterium* cultures were first diluted from an overnight culture to an OD_600_ of 0.05 in BHI and grown until an OD_600_ of 0.5 was reached. Serial dilutions were performed up to 10^-10^, and 150 μL of each dilution was inoculated onto BHI plates containing increasing erythromycin concentrations (0.12-4 mg/L). Plates were incubated for 24-48 hours, and CFU/mL was determined by colony counting; percent survival was calculated.

### *Drosophila* oral infection assay

Flies were maintained in standard culture conditions in media containing sugar, molasses, and yeast and kept at room temperature. Three independent experimental replicates were performed, each using 20 adult wild-type *Drosophila melanogaster* flies (*w^1118^* strain). We followed an oral infection protocol previously reported (15,16). For each assay, 5 mL of overnight bacterial culture of the test strains (JVH47 and P4526) was prepared, together with a non-inoculated control and *Pseudomonas aeruginosa* PAO1 as the positive control. Cultures were centrifuged at 4,000 × g for 10 min, and the resulting pellets were resuspended in fresh BHI medium and adjusted to an OD_600_ of 0.5. Bacterial suspensions were centrifuged again, and the pellets were resuspended in 500 µL of a 50 mM sucrose solution.

For fly exposure, 100 µL of the bacterial suspension was applied as the upper layer onto a sucrose-agar feeding medium (5% sucrose, 2% agar). 20 adult flies pre-fed on a 50 mM sucrose solution for 48 h were transferred to each vial. Experiments were conducted at room temperature, and fly survival was monitored by counting living individuals usually at 24-h intervals until the end of the experiment.

Fly survival data were analyzed using Kaplan–Meier survival curves. All experiments were performed in triplicate, and data from independent replicates were pooled for analysis. Statistical significance was considered at p < 0.05.

### Cell culture and cell viability assay

Caco-2 cells (ATCC HTB-37 ™) were cultured in DMEM containing 10% fetal bovine serum (FBS), 5 µg/ml penicillin-streptomycin, and 5 mM sodium pyruvate and incubated at 37 °C in a humidified atmosphere with 5% CO2. Caco-2 cells (1 × 10^4^) were seeded in 96-well plates and cultured for 3 days post-confluence for the cell viability assay. Cells were treated with cell-free supernatants for 24 h. Cell viability was determined by 3-(4,5-dimethylthiazol-2-yl)–2,5-diphenyltetrazolium bromide (MTT) assay (Sigma-Aldrich, St. Louis, MO, USA). Cells were incubated with 0.5 mg/ml MTT reagent at 37 °C for 4 h. Acidified 20% SDS was used for 30 min to resuspend the purple crystals. SpectraMax iD3 Multi-Mode Microplate Reader measured the optical density at 570 nm. The experiments were repeated in triplicate across at least four independent experiments, and the results were expressed as the mean percentage of viable cells and compared with the control (untreated). Survival analyses and graphical representations were generated using GraphPad Prism.

### Genome sequencing and annotation

The *Exiguobacterium* strains JVH47 and P4526 were incubated at 37 °C in LB medium at 150 rpm for 24 hours. Genomic DNA was extracted using the phenol-chloroform method and sent for sequencing at SeqCenter (USA). Genomic DNA was used to construct sequencing libraries using the Illumina DNA Prep kit and IDT 10 bp UDI indices, following Illumina’s recommended protocols. Libraries were sequenced on an Illumina NextSeq 2000, producing 2×151-bp reads. Demultiplexing, quality control, and adapter trimming were performed using Illumina’s bcl-convert v3.9.3, yielding 91.381% of base pairs with> Q30 quality. Poor-quality readings were filtered using Trimmomatic v0.36 software [13]. Readings with sequencing quality scores above 28 and lengths exceeding 36 base pairs were used for the de novo assembly of Illumina reads with SPAdes v3.13.1[14], using k-mer sizes of 21, 33, 55, and 77. The quality of the genome assembly was assessed using QUAST v5.2.0 [15]. The Prokka v1.14.6 software [16] was used to predict the open reading frames (ORFs) of the assembled genome using default parameters for Gram-positive bacteria.

### Pangenome reconstruction, functional annotation, and gene content variation analysis

Genomes of the strains JVH47 and P4526 were compared against the other 146 genomes of *Exiguobacterium* strains. Genomes were selected based on 16S homology; genomes with completeness above 96% were retrieved from GenBank (Supplementary data 1). The annotated genome assemblies were used as input to a pangenome analysis with the Panaroo pipeline [17]. Orthologous genes for all ORFs across the 148 genomes were grouped using CD-HIT [18], with a protein sequence identity threshold of at least 90%; each cluster contained only a single paralog per genome. Potentially contaminating ORFs were removed from the dataset. ORFs present in ≥99% of strains were considered as core genes, genes classified as soft core were those present in the range of ≥95% and <99% genomes, genes present between ≥15% and <95% of strains were classified as shell genes, and cloud genes were those present in less than 15% of *Exiguobacterium* genomes.

Alignment of core genes was generated with MAFFT [18], and a maximum-likelihood phylogenetic tree was constructed from this alignment using IQ-TREE v2.4.0 [19], with ultrafast bootstrap [20] applied with 1000 replicates. ModelFinder [21] determined that the phylogenetic tree was best fitted to the model GTR+F+I+R9. iTOL v7.4.2 facilitated visualization of the phylogenetic tree.

Additionally, predicted ORFs were use for functional annotation based on gene orthology using blastKOALA v3.1 [22] from the Kyoto Encyclopedia of Genes and Genomes (KEGG) database, available at https://www.kegg.jp/blastkoala/. Resulted KEGG orthology (KO) entries were further related to the Clusters of Orthologous Genes (COG) database using the BRITE binary relation files (available at https://www.kegg.jp/brite/br08906). Thus, each KO entry was classified into COG categories. A gene could be assigned to more than one functional COG category.

### Detection of mobile genetic elements (MGEs), defense mechanism systems, virulence factors and ARGs

ABRICATE (https://github.com/tseemann/abricate) was used to screen the assembly sequences against the NCBI AMRFinderPlus {Feldgarden, 2021 #34}, CARD [23], ARG-ANNOT [24] MEGARES 3.00 [25] and PlasmidFinder [26] databases for the detection of antimicrobial resistance genes (ARGs). Information of all databases was concatenated, redundant ORFs were removed, and the NCBI annotation was conserved when the same ORF was annotated differently across databases (Supplementary data 1).

Additionally, sequences of plasmids and viruses were screened using geNomad v1.11.1 [27] under conservative filtering. geNomad was run on the Galaxy server available at https://usegalaxy.eu/ [28]

Databases of genes responding to UV and osmotic stress (Supplementary data 1) were constructed based on information reported for bacteria [29] [30] [31] [32] [33] [34] [35]. BLAST was used to align amino acid sequences from both databases against the annotated genome assemblies of strains JVH47 and P4526, and against the clustered ORFs of the previously analyzed Exiguobacterium genomes obtained from CD-HIT. Sequences with at least 30% sequence identity were classified as either UV- or osmotic-stress genes.

Biosynthetic gene cluster prediction was performed using antiSMASH v8.0.4 [36].

### UV exposure assay

Bacterial UV resistance was evaluated using a UV-C survival assay. Five-day-old cultures were grown in LB medium at 30 °C, then washed and resuspended in sterile phosphate-buffered saline (PBS). Cell suspensions were standardized to an OD_600_ of 0.1. Aliquots of 100 µL were spread as a thin, uniform layer onto sterile Petri dishes and exposed to UV-C radiation (254 nm), leaving a distance of 10 cm from the lamp. Samples were irradiated for up to 3 h, whereas non-irradiated controls were protected from light using aluminum foil. Immediately after exposure, cells were recovered in PBS, serially diluted, and plated on LB agar. Plates were incubated at 30 °C in the dark to prevent photoreactivation, and colony-forming units (CFUs) were counted after 24–48 h. Survival was calculated as the ratio of CFUs from irradiated samples to non-irradiated controls.

### Carotenoid extraction and quantification

Carotenoid production was assessed in bacterial cultures by comparing production in solid and liquid LB media. In both cases, a pre-culture in LB medium was adjusted to an OD_600_ of 0.1, an aliquot of 100 µL was spread onto LB agar, and 300 µL was inoculated in 30 mL of LB medium. In both cases, bacteria were at 30 °C for 5 days. Cells from a Petri dish were collected in centrifuge tubes and resuspended in PBS; cells from liquid culture were harvested by centrifugation. In each case, cells were washed once with sterile PBS. Pigments were extracted by resuspending the cell pellets in methanol and incubating at 60 °C for 20–30 min with occasional mixing. Cell debris was removed by centrifugation, and the absorbance of the supernatant was measured spectrophotometrically at 450 nm. Total carotenoid concentration was estimated as β-carotene equivalents using the equation: “ *Carotenoids* (μ*g*/*mL*)” = (*A*_450 x *V* x 10^4)/(*A*_1“ *cm*” ^(1%) x *l*),where *A*_450is the absorbance at 450 nm, Vis the extraction volume (mL), *l* is the path length of the cuvette (cm), and *A*_1“ *cm*” ^(1%) = 2592corresponds to the specific absorption coefficient of β-carotene in methanol. Carotenoid content was normalized to wet cell biomass (µg β-carotene equivalents mg⁻¹ wet weight).

### Data availability

The genomes of strains JVH47 and P4526 were deposited in GenBank under BioProject accession numbers PRJNA1415846 and PRJNA1415850, respectively.

## Results

### Isolation and identification of *Exiguobacterium* strains

Following an urban flooding event in Mexico City, we isolated a morphologically diverse bacterial community from sediment samples containing a mixture of pluvial and residual water. One isolate, found at a high frequency (48.4%), grew on a selective medium containing a mixture of antibiotics. This isolate was selected for further characterization. The colony displayed a distinctive orange pigmentation and a smooth, shiny, circular morphology with well-defined edges. Gram staining revealed a Gram-positive, short rod-shaped bacterium.

Amplification and sequencing of the 16S rRNA gene indicated that the isolate shared 96.48% sequence similarity with *Exiguobacterium mexicanum* strain [accession number OK1358331], and 96.36% similarity with *Exiguobacterium aquaticum* strain [accession number MF5922911]. We designated this isolate as *Exiguobacterium* sp. JVH47.

To contextualize the adaptation of JVH47, which originated from a polluted urban environment, we compared it to an *Exiguobacterium* strain isolated from a less anthropogenically impacted habitat. The comparison strain, P4526, was obtained from the Cuatro Ciénegas Basin in Mexico, an ecologically diverse region comprising aquatic systems ranging from ephemeral ponds to permanent lakes, rivers, and springs [37].

### Genome Assembly

After confirming that neither sample had been previously reported, the genomes of Exiguobacterium strains JVH47 and P4526 were sequenced using Illumina technology and assembled de novo with SPAdes. For JVH47, the assembly yielded 49 contigs (≥500 bp), resulting in a total genome size of 2.97 Mb and a GC content of 51.97%. The largest contig was 633,504 bp, with an N50 of 130,672 bp and an L50 of 5. In contrast, the P4526 assembly produced 22 contigs (≥500 bp), totaling 2.87 Mb with a slightly higher GC content of 52.46%. The assembly for P4526 was more contiguous, with the largest contig of 1.47 Mb, an N50 of 1,467,817 bp, and an L50 of 1 (Table S1). These statistics demonstrated that the assembled genomes of JVH47 and P4526 were not highly fragmented and were suitable for comparison with other available *Exiguobacterium* genomes.

### Genome Annotation and Gene Ontology Analysis

According to the COG classification, approximately 7% of predicted genes in both strains are involved in translation and biogenesis. The second and third more abundant classes of genes are involved in amino acid and carbon transport and metabolism, each representing around 5-6% of predicted ORF. Notably, the function of approximately half of the predicted genes remain unknown (Fig. S1), underscoring the paucity of information about the genus.

### Genome comparison

Using the genomic context of strains JVH47 and P4526, we compared their genomes with 146 reported *Exiguobacterium* genomes. Selected genomes spanned sizes from 2.60 Mb (*Exiguobacterium algae* strain S126) to 3.36 Mb (*Exiguobacterium* sp. s191) and GC content between 47.09% and 54.99%. Panaroo estimated a pan-genome size of 19,593 clustered genes. In accordance with conservation levels (refer to methodology section), the analysis showed that the core genome comprised 76 genes (0.4%), 6,533 (33.3%) were shell genes, and 12,984 were categorized as cloud genes, representing 66.3% of the pan-genome. A total of 5,507 genes (28.1%) were present in only one genome. While the majority of core genes were assigned to the functions “translation, ribosomal structure and biogenesis”, a notable proportion of cloud genes with known functions are associated with carbohydrate transport and metabolism (Fig. S1). The genomes of strains *Exiguobacterium* sp. s192 and *Exiguobacterium undae* strain KCTC 3810 had the largest cloud of genes sampled to date within the genus, accounting for 926 and 874 genes, respectively. The strain JVH47 contained 223 cloud genes, and the strain P4526 contained 235 cloud genes; in both cases, most annotated genes were involved in carbohydrate transport and metabolism. To further compare the genomes of *Exiguobacterium* strains, we constructed a maximum-likelihood phylogenetic tree (Fig. 1) based on an alignment of core genes. The phylogeny clustered *Exiguobacterium* strains into two main groups with high confidence, with most branches displaying bootstrap values above 90%. A major group consists of 93 evolutionarily related strains that harbor JVH47 and P4526. JVH47 is closely related to the strains s26 and s151, which were also sourced from soil sediment in China. On the other hand, P4526 was phylogenetically related to other strains isolated from various sources, mainly from soil sediments. These results altogether highlight the high variability in gene content among strains of the genus *Exiguobacterium*.

**Fig 1.**
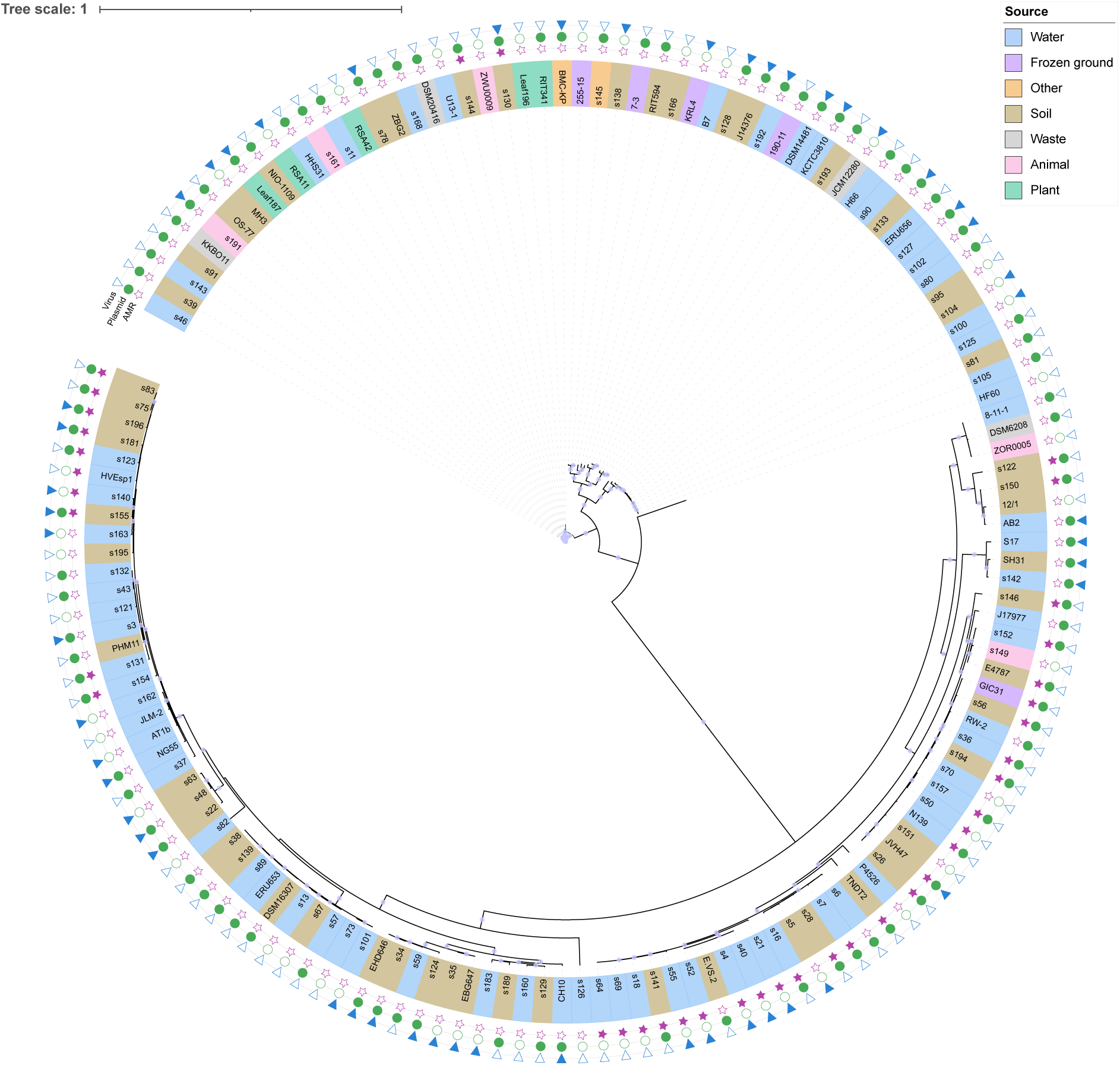
Phylogenetic tree of the genus *Exiguobacterium*. Branches with bootstrap value above 90 display a circle in the middle of the branch. The ID of each strain is coloured according to the isolation source. Strains harbouring genes conferring antibiotic resistance are shown with a fully-coloured purple star, strains with plasmid sequences are highlighted with a filled green circle and strains containing virus sequences are depicted with a filled blue triangle.

### Phenotypic Characterization of *Exiguobacterium* sp. strains

Strains JVH47 and P4526 exhibited broad environmental tolerance, demonstrating the ability to grow under a wide range of pH, salinity, and temperature conditions (Fig. 2). Both strains were alkali-resistant, maintaining growth at pH values up to 11, but neither could grow in acidic environments below pH 6 (Fig. 2a). In salinity assays, both strains showed moderate halotolerance, with sustained growth observed at NaCl concentrations up to 5% (Fig. 2b), only discreate growth was observed at NaCl 10 %. Both strains contain more than 30 genes reported to confer osmotic tolerance (Table S2), including genes involved in potassium import and genes that produce osmo-protectant solutes, such as glutamate or proline [33–35]. Temperature tolerance tests revealed no growth at 4 °C for either strain (Fig. 2c). However, both strains demonstrated thermotolerance, sustaining growth at 40 °C.

**Fig 2.**
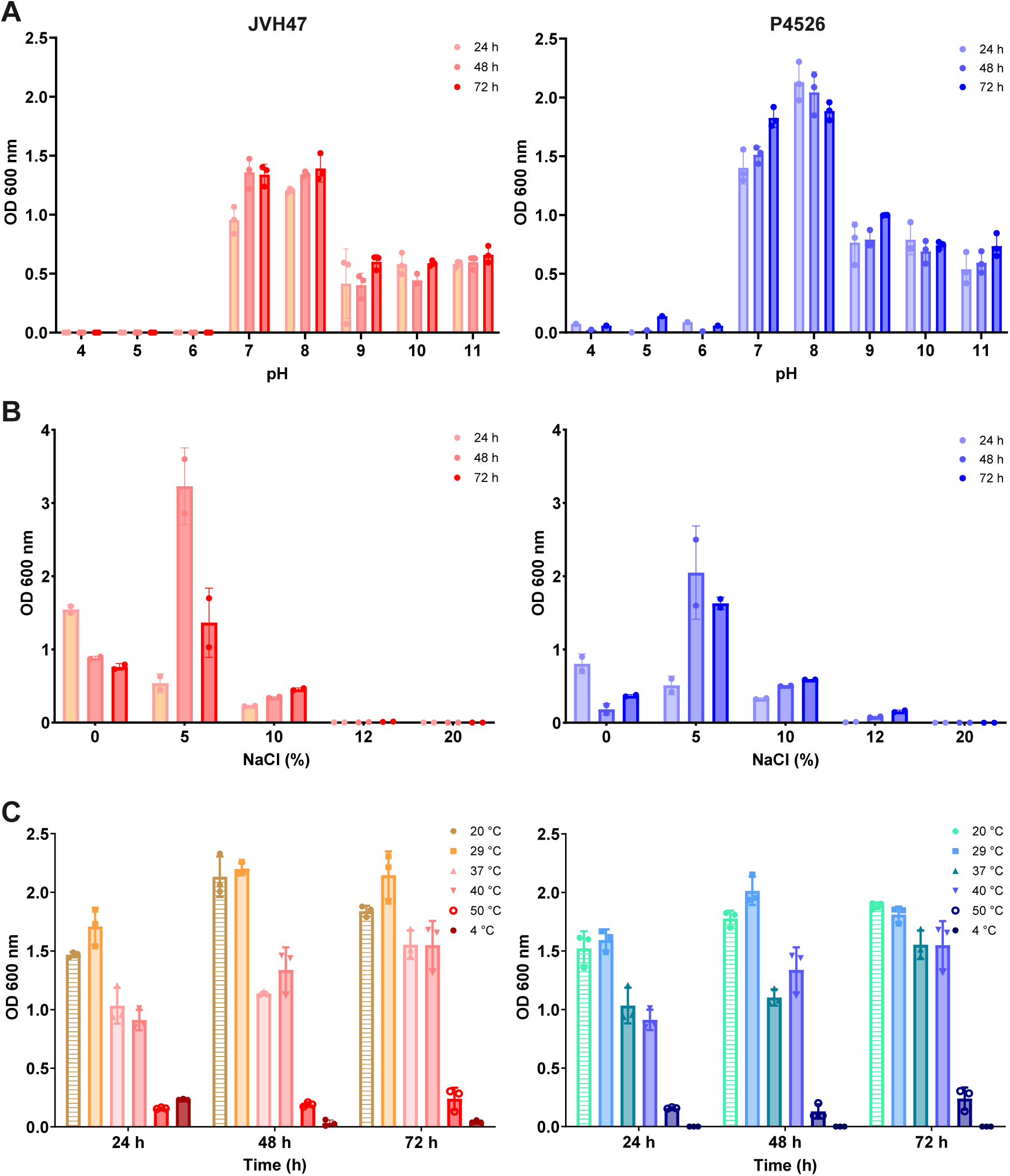
Comparative evaluation of tolerance to extreme conditions in *Exiguobacterium* strains JVH47 and P4526. **a)** Growth analysis under different pH. **b)** Halotolerance assessment. **c)** Growth evaluation at different temperatures. Results for the strain JVH47 are depicted in a gradient of pink-red on the left, whereas that results for the strain P4526 are shown on the right side in a green-blue gradient.

### UV resistance

Several *Exiguobacterium* strains have been isolated from environments characterized by high ultraviolet (UV) radiation. For example, recently the resistance of *Exiguobacterium* sp. S17—isolated from modern stromatolites—to UV-B radiation and identified a comprehensive UV-resistome (UVres) was demonstrated [38]. This UVres encompasses multiple functional subsystems, including UV sensing and response regulation, avoidance and shielding mechanisms, DNA damage repair pathways, oxidative stress responses, energy management, and metabolic resetting. Inspired by these findings, we evaluated the UV resistance of strains JVH47 and P4526 to determine their capacity to withstand UV-induced stress and to explore whether environmental origin correlates with UV resilience. In both strains, viable cells were recovered after 2 and 3 h of UV irradiation (Fig. 3a). Previous work has demonstrated that pigment production is associated with adaptation to extreme environments and UV protection [39]. Consequently, partial carotenoid production was evaluated. As shown, strain P4526 exhibited significantly higher carotenoid production than JVH47 (Fig. 3b).

**Fig 3.**
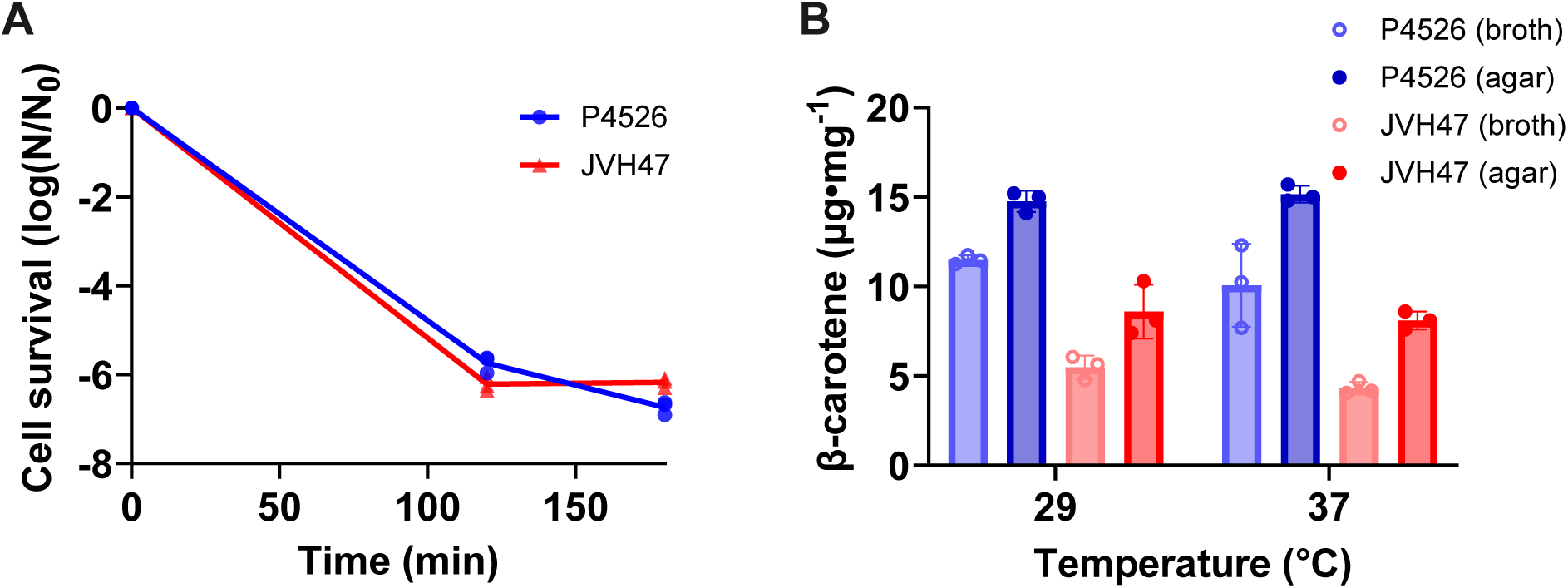
Evaluation of resilience to UV irradiation in *Exiguobacterium* strains JVH47 and P4526. **a)** Growth evaluation after exposure to UV for up to 3 h. Survival was calculated as the ratio of colony forming units from irradiated samples to non-irradiated cells. **b)** Comparison of carotenoid production in strains JVH47 and P4526 in liquid and solid LB media, production was evaluated at 29 °C and 37 °C.

We analyzed the genomes of both strains to identify genes conferring UV resistance. The *in silico* analysis revealed that genes associated with the UV resistome were present in both genomes, these genes accounted for 2% of the total ORFs in each genome (Table S2). Additionally, we examined genes related to carotenoid biosynthesis. Biosynthetic gene cluster prediction in each strain using antiSMASH v8.0.4 [36] anticipated two terpene-like clusters (Fig. S2), and both genomes carry a geranylgeranyl diphosphate synthase (Supplementary data 1), which encodes a protein reported to catalyze the first reaction of carotenoid biosynthesis in tomato [40] These genes may be responsible for the carotenoid production. Therefore, the observed differences in UV tolerance may be attributed, at least in part, to differential gene expression and/or variation in carotenoid composition.

### Virulence factor and *in vitro* pathogenicity

*Exiguobacterium* are mostly environmental bacteria, but some reports of clinical disease have implicated these bacteria [9, 11]. To assess potential virulence and cytotoxic effects, two approaches were considered: infection in *D. melanogaster* and cytotoxicity in Caco-2 cells.

Taking advantage of *D. melanogaster* as a reliable model for host–pathogen interactions [41], the virulence of the *Exiguobacterium* strains was evaluated alongside the well-characterized pathogen *P. aeruginosa* [42]. As shown, fly survival profiles varied depending on the bacterial strain used for oral exposure (Fig. 4a). As expected, flies exposed to *P. aeruginosa* PAO14 showed a clear trend toward reduced survival over time, with mortality initiating earlier compared to the other treatments. In contrast, flies fed with strain JVH47 tended to maintain higher survival probabilities throughout the experiment, largely overlapping with or slightly exceeding those observed in the non-inoculated control. Exposure to strain P4526 resulted in an intermediate survival pattern, with a gradual decline in survival that appeared more pronounced than the control but less severe than that observed for PAO14.

**Fig 4.**
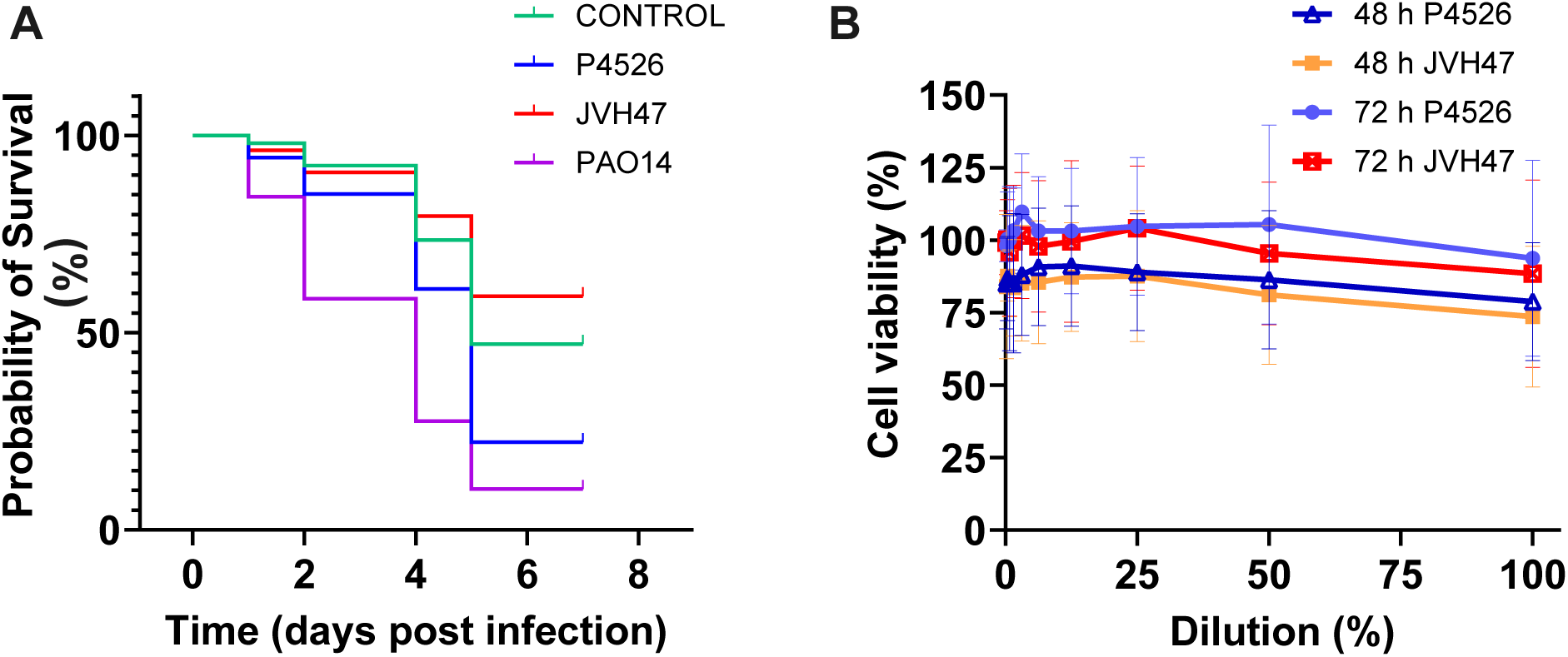
*In vitro* evaluation of toxicity mediated by *Exiguobacterium* strains JVH47 and P4526. **a)** Evaluation of cell survival of Drosophila melanogaster to oral infection with strains JVH47 and P4526 and the positive control *Pseudomonas aeruginosa* PAO1. **b)** Evaluation of viability of Caco-2 cells treated with supernatants of strains JVH47 and P4526.

To further assess the cytotoxic potential of *Exiguobacterium* strains, we performed MTT assays on Caco-2 cells using serially diluted cell-free supernatants (CFS) of JVH47 and P4526. CFS were collected from JVH47 and P4526 strains of 48 and 72 h. After 24 h of exposure to CFS, no reduced cell viability was observed (Fig. 4b).

### Antimicrobial Resistance Genes and Mobile Genetic Elements in *Exiguobacterium* **Genomes**

At the conditions tested, only the strain P4526 displayed attenuated pathogenicity against *D. melanogaster*. However, *Exiguobacterium* strains could be potential pathogens [9]. To explore the pathogenicity potential of the genus, we surveyed their genomes for mobile genetic elements –including mobilome genes and plasmid sequences, competence-related genes, conjugation genes, virus sequences, and antimicrobial resistance genes. Overall, 35 strains possess the gene *insK*, a putative transposase used for the insertion of the sequence element IS150. The strain *Exiguobacterium* sp. KRL4 contains the gene ltrA, which encodes a group II intron-encoded protein (Supplementary data 1). Additionally, 51 strains contain a virus sequence in their genomes, and 97 strains contain at least one plasmid element (Fig. 1), with MOBP2 HMM (Hidden Markov Model) profile of the plasmid MobL relaxases family as the most prevalent (Supplementary data 1). The MOBP2 HMM profile has been identified in mobilizable plasmids [43] which means that require a helper conjugative plasmid to be transmissible [44]. Although mostly alone, MOBP2 HMM profile could be found altogether with the relaxases MOBP1, MOBP3, MOBV and/or T_virB11. The latest has been described as part of the virulence regulon *virB* in *Agrobacterium tumefaciens* and could assist in DNA transfer [45]. This virulence-associated gene was found in the genomes of Exiguobacterium sibiricum 7-3 and *Exiguobacterium* sp. KKBO11, *Exiguobacterium* sp. RIT594 and *Exiguobacterium* sp. s90, from which the strains 7-3 and RIT594 are closely phylogenetically related (Fig. 1). Furthermore, *Exiguobacterium* sp. S17 contained one T4CP2, a type IV coupling protein of the type IV secretion system, which is involved in horizontal gene transfer [46].

The strain JVH47 harbors numerous mobile elements, as evidenced by COG annotation, including one transposase belonging to the IS30 family and 9 putative transposases. The strain also possesses a single MOBP2 (Table S2). Likewise, the strain P4526 contains 5 putative transposases. Both strains contain members of the *comE, comF* and *comG* operons, including the gene encoding the competence protein ComGC, a principal component of the type IV-like pilus [47] and the gene *comFA*, which has been found in all naturally competent Gram-positive bacteria [48–50]. Even when the strain JVH47 carries more genes involved in pathogenicity, their expression could be tightly regulated, which may account for the lack of *in vitro* pathogenicity.

From the 148 *Exiguobacterium* strains that constitute our pan-genome, only 39 strains possess at least one antimicrobial resistance gene. The strain *Exiguobacterium* sp. s150, which was sourced from soil in a pig farm in China, contains 9 genes that confer resistance to different classes of antibiotics – the highest number of resistance genes in a single strain – including resistance to aminoglycoside, trimethoprim, macrolide, bleomycin, tetracycline and lincosamide antibiotics (Fig. 5a). Additionally, the strain harbors the multiresistance plasmid pKKS825 that contribute to resistance to aminoglycosides, tetracyclines and trimethoprim [51].

**Fig 5.**
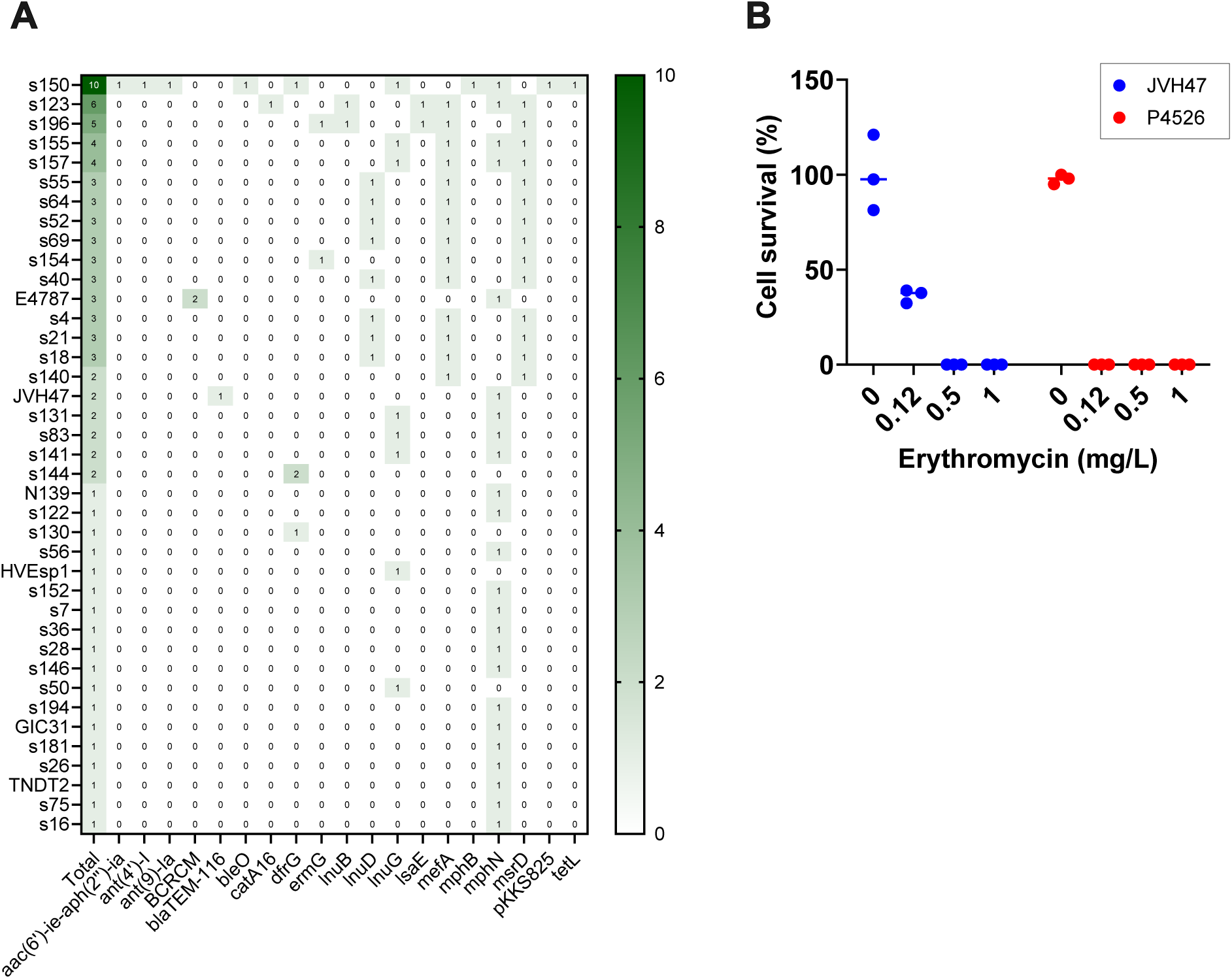
Survey of antimicrobial resistance in *Exiguobacterium* strains. **a)** Genes conferring antibiotic resistance located in the genomes of *Exiguobacterium* strains. **b)** Survival evaluation of strains JVH47 and P4526 under increasing concentrations of erythromycin.

Among the resistance phenotypes identified, macrolide resistance was the most prevalent, with three macrolide resistance genes identified (Fig. 5a). The gene *mphN*, encoding a macrolide 2’-phosphotransferase, was found in 24 of 39 strains that carry antimicrobial resistance genes, whereas *mefA* and *msrD* were found in the same 14 strains each. Interestingly, *mefA* was located immediately upstream of msrD, consistent with an operon architecture. In *Streptococcus pneumoniae*, both MefA and MsrD associates forming an efflux complex to eject macrolides [52] Other genes conferring resistance to macrolides present in *Exiguobacterium* strains at lower frequencies were mphB and ermG, which encode a methyltransferase. The presence of these genes highlights that the resistance to macrolides in *Exiguobacterium* bacteria is mainly conferred through two mechanisms: efflux-mediated (*mefA* and *msrD*) and chemical modification of the antibiotic (*mphN*, *mphB* and *ermG*).

### Erythromycin Resistance and analysis of *mphN*

Macrolide resistance is widespread among pathogenic bacteria and is typically caused by GTP-dependent macrolide kinases (Mph), ribosomal methyltransferases, or efflux pumps. Mph enzymes confer resistance by phosphorylating the 2′-OH group of the critical dimethylamine sugar in macrolides, which blocks their binding to the ribosome [53]. We ask whether the presence of an *mphN* gene confers erythromycin resistance in the original host. To this end, we compared the average cell survival in the presence of erythromycin (Ery) between JVH47 and P4526. Average cell survival at the indicated Ery concentrations was normalized to the survival percentage of the culture grown without antibiotics. Both strains were sensitive to Ery; however, JVH47 exhibited a partial change in antibiotic susceptibility. At an erythromycin concentration of 0.12 mg/L, JVH47 showed an average cell survival of approximately 40%, whereas no survival was detected for P4526 under the tested antibiotic concentrations (Fig. 5b). However, after several passages, the resistance observed was lost. This phenotype is consistent with the genetic information: the JVH47 strain harbors the *mphN* gene, whereas P4526 lacks any resistance gene.

## Discussion

The genus *Exiguobacterium* is gaining relevance due to its predominance in environmental samples [54–56], and yet, there is still little information about the genus. Most Exiguobacterium strains isolated to date have been obtained from extreme environments, highlighting their adaptability and potential for biotechnological applications. Nevertheless, there are reports of *Exiguobacterium* strains forming biofilm [57] and even strains causing infections to human [9] and animals [58]. Thus, it is important to unveil the factors influencing the *Exiguobacterium* strains become pathogenic.

In this study, we describe the *Exiguobacterium* strains JVH47 and P4526, with special emphasis in understanding if the habitat influences the acquisition of virulence and AMR genes. For this, we tested the adaptation capabilities of both strains and performed a comprehensive analysis of the pathogenicity of these two novel strains and the other 146 previously reported strains.

Our results are consistent with previous studies on the genus *Exiguobacterium*. However, we detected a different grouping pattern when comparing our phylogenetic tree with a phylogeny previously reported by Zheng’s group in 2021[1]. The observed differences may reflect variations in methodological approaches. With the methodology applied in our work, we obtained a larger pan-genome size (19,593 gene clusters against 8,857 previously reported), nevertheless, most of the predicted ORFs are hypothetical proteins.

Our genomic analysis demonstrated that strains in the genus possess the genetic arsenal required to overcome osmotic stress and UV damage, enabling halotolerance and recovery from UV damage. We also found that some strains possess virulence genes. Comparing JVH47, which can be classified as an “urban” strain, with the strain P4526, which was not exposed to human activity, we observed differences in the abundance of mobile elements, with JVH47 containing more, possibly because the strain acquired them during its cosmopolitan distribution. Nevertheless, JVH47 did not displayed a mortal pathogenicity phenotype.

## Conclusion

The study of *Exiguobacterium* and other highly adaptable bacterial genera is critical for the early identification of emerging pathogens and for preventing their escalation into serious threats to human health. Heavily impacted and polluted environments impose intense selective pressures that actively drive the acquisition and dissemination of antimicrobial resistance and virulence genes. These environments function as evolutionary hotspots, facilitating bacterial adaptation with direct consequences for public health. These findings highlight an urgent need for systematic environmental surveillance and immediate actions to improve environmental conditions, mitigate microbial risks, and safeguard human, animal, and ecosystem health within a One Health framework.

## Acknowledgments

Authors acknowledge Dr. Valeria Souza for generously providing *Exiguobacterium* sp. P4526 used in this study.

## Funding and additional information

This work was partially supported by grants from UNAM-DGAPA PAPIIT IA206823 (A.R.R.)

## References

1. Zhang D, Zhu Z, Li Y, Li X, Guan Z, Zheng J. Comparative Genomics of Exiguobacterium Reveals What Makes a Cosmopolitan Bacterium. mSystems. 2021;6(4):e00383–21. doi: 10.1128/mSystems.00383-21.

2. Yang Y, Yang J, Wu W-M, Zhao J, Song Y, Gao L, et al. Biodegradation and Mineralization of Polystyrene by Plastic-Eating Mealworms: Part 2. Role of Gut Microorganisms. Environmental Science & Technology. 2015;49(20):12087–93. doi: 10.1021/acs.est.5b02663.

3. Orozco-Medina C, López-Cortés A, Maeda-Martínez AM. Aerobic Gram-positive heterotrophic bacteria Exiguobacterium mexicanum and Microbacterium sp. in the gut lumen of Artemia franciscana larvae under gnotobiotic conditions. Current Science. 2009;96(1):120–9.

4. Hossain TJ, Chowdhury SI, Mozumder HA, Chowdhury MNA, Ali F, Rahman N, et al. Hydrolytic Exoenzymes Produced by Bacteria Isolated and Identified From the Gastrointestinal Tract of Bombay Duck. Frontiers in microbiology. 2020;11:2097. Epub 2020/09/29. doi: 10.3389/fmicb.2020.02097. PubMed PMID: 32983064; PubMed Central PMCID: PMCPMC7479992.

5. Kasana RC, Pandey CB. Exiguobacterium: an overview of a versatile genus with potential in industry and agriculture. Critical Reviews in Biotechnology. 2018;38(1):141–56. doi: 10.1080/07388551.2017.1312273.

6. Cong M, Jiang Q, Xu X, Huang L, Su Y, Yan Q. The complete genome sequence of Exiguobacterium arabatum W-01 reveals potential probiotic functions. MicrobiologyOpen. 2017;6(5). Epub 2017/06/08. doi: 10.1002/mbo3.496. PubMed PMID: 28589562; PubMed Central PMCID: PMCPMC5635162.

7. Hadi JA, Gutierrez N, Alfaro AC, Roberts RD. Use of probiotic bacteria to improve growth and survivability of farmed New Zealand abalone (Haliotis iris). New Zealand Journal of Marine and Freshwater Research. 2014;48(3):405–15. doi: 10.1080/00288330.2014.909857.

8. Kim S, Jeon H, Bai SC, Hur J-W, Han H-S. Evaluation of Salipiger thiooxidans and Exiguobacterium aestuarii from the Saemangeum Reservoir as Potential Probiotics for Pacific White Shrimp (Litopenaeus vannamei). Microorganisms [Internet]. 2022; 10(6).

9. Chen X, Wang L, Zhou J, Wu H, Li D, Cui Y, et al. Exiguobacterium sp. A1b/GX59 isolated from a patient with community-acquired pneumonia and bacteremia: genomic characterization and literature review. BMC infectious diseases. 2017;17(1):508. Epub 2017/07/25. doi: 10.1186/s12879-017-2616-1. PubMed PMID: 28732529; PubMed Central PMCID: PMCPMC5521131.

10. Keynan Y, Weber G, Sprecher HJJomm. Molecular identification of Exiguobacterium acetylicum as the aetiological agent of bacteraemia. 2007;56(4):563–4.

11. Kenny F, Xu J, Millar B, McClurg R, Moore JJBjobs. Potential misidentification of a new Exiguobacterium sp. as Oerskovia xanthineolytica isolated from blood culture. 2006;63(2):86.

12. Pitt TL, Malnick H, Shah J, Chattaway MA, Keys CJ, Cooke FJ, et al. Characterisation of Exiguobacterium aurantiacum isolates from blood cultures of six patients. Clinical Microbiology and Infection. 2007;13(9):946–8. doi: 10.1111/j.1469-0691.2007.01779.x.

13. Bolger AM, Lohse M, Usadel B. Trimmomatic: a flexible trimmer for Illumina sequence data. Bioinformatics. 2014;30(15):2114–20. doi: 10.1093/bioinformatics/btu170 %J Bioinformatics.

14. Prjibelski A, Antipov D, Meleshko D, Lapidus A, Korobeynikov A. Using SPAdes De Novo Assembler. Current protocols in bioinformatics. 2020;70(1):e102. Epub 2020/06/20. doi: 10.1002/cpbi.102. PubMed PMID: 32559359.

15. Mikheenko A, Prjibelski A, Saveliev V, Antipov D, Gurevich A. Versatile genome assembly evaluation with QUAST-LG. Bioinformatics. 2018;34(13):i142–i50. Epub 2018/06/29. doi: 10.1093/bioinformatics/bty266. PubMed PMID: 29949969; PubMed Central PMCID: PMCPMC6022658.

16. Seemann T. Prokka: rapid prokaryotic genome annotation. Bioinformatics. 2014;30(14):2068–9. Epub 2014/03/20. doi: 10.1093/bioinformatics/btu153. PubMed PMID: 24642063.

17. Tonkin-Hill G, MacAlasdair N, Ruis C, Weimann A, Horesh G, Lees JA, et al. Producing polished prokaryotic pangenomes with the Panaroo pipeline. Genome Biology. 2020;21(1):180. doi: 10.1186/s13059-020-02090-4.

18. Katoh K, Misawa K, Kuma Ki, Miyata T. MAFFT: a novel method for rapid multiple sequence alignment based on fast Fourier transform. Nucleic Acids Research. 2002;30(14):3059–66. doi: 10.1093/nar/gkf436 %J Nucleic Acids Research.

19. Minh BQ, Schmidt HA, Chernomor O, Schrempf D, Woodhams MD, von Haeseler A, et al. IQ-TREE 2: New Models and Efficient Methods for Phylogenetic Inference in the Genomic Era. Mol Biol Evol. 2020;37(5):1530–4. Epub 2020/02/06. doi: 10.1093/molbev/msaa015. PubMed PMID: 32011700; PubMed Central PMCID: PMCPMC7182206.

20. Hoang DT, Chernomor O, von Haeseler A, Minh BQ, Vinh LS. UFBoot2: Improving the Ultrafast Bootstrap Approximation. Mol Biol Evol. 2018;35(2):518–22. Epub 2017/10/28. doi: 10.1093/molbev/msx281. PubMed PMID: 29077904; PubMed Central PMCID: PMCPMC5850222.

21. Kalyaanamoorthy S, Minh BQ, Wong TKF, von Haeseler A, Jermiin LS. ModelFinder: fast model selection for accurate phylogenetic estimates. Nature Methods. 2017;14(6):587–9. doi: 10.1038/nmeth.4285.

22. Kanehisa M, Sato Y, Morishima K. BlastKOALA and GhostKOALA: KEGG Tools for Functional Characterization of Genome and Metagenome Sequences. Journal of molecular biology. 2016;428(4):726–31. Epub 2015/11/21. doi: 10.1016/j.jmb.2015.11.006. PubMed PMID: 26585406.

23. Alcock BP, Huynh W, Chalil R, Smith KW, Raphenya AR, Wlodarski MA, et al. CARD 2023: expanded curation, support for machine learning, and resistome prediction at the Comprehensive Antibiotic Resistance Database. Nucleic Acids Res. 2023;51(D1):D690-d9. Epub 2022/10/21. doi: 10.1093/nar/gkac920. PubMed PMID: 36263822; PubMed Central PMCID: PMCPMC9825576.

24. Gupta SK, Padmanabhan BR, Diene SM, Lopez-Rojas R, Kempf M, Landraud L, et al. ARG-ANNOT, a new bioinformatic tool to discover antibiotic resistance genes in bacterial genomes. Antimicrobial agents and chemotherapy. 2014;58(1):212–20. Epub 2013/10/23. doi: 10.1128/aac.01310-13. PubMed PMID: 24145532; PubMed Central PMCID: PMCPMC3910750.

25. Bonin N, Doster E, Worley H, Pinnell LJ, Bravo JE, Ferm P, et al. MEGARes and AMR++, v3.0: an updated comprehensive database of antimicrobial resistance determinants and an improved software pipeline for classification using high-throughput sequencing. Nucleic Acids Res. 2023;51(D1):D744–d52. Epub 2022/11/17. doi: 10.1093/nar/gkac1047. PubMed PMID: 36382407; PubMed Central PMCID: PMCPMC9825433.

26. Carattoli A, Zankari E, García-Fernández A, Voldby Larsen M, Lund O, Villa L, et al. In silico detection and typing of plasmids using PlasmidFinder and plasmid multilocus sequence typing. Antimicrobial agents and chemotherapy. 2014;58(7):3895–903. Epub 2014/04/30. doi: 10.1128/aac.02412-14. PubMed PMID: 24777092; PubMed Central PMCID: PMCPMC4068535.

27. Camargo AP, Roux S, Schulz F, Babinski M, Xu Y, Hu B, et al. Identification of mobile genetic elements with geNomad. Nature biotechnology. 2024;42(8):1303–12. Epub 2023/09/22. doi: 10.1038/s41587-023-01953-y. PubMed PMID: 37735266; PubMed Central PMCID: PMCPMC11324519.

28. Community TG. The Galaxy platform for accessible, reproducible, and collaborative data analyses: 2024 update. Nucleic Acids Research. 2024;52(W1):W83–W94. doi: 10.1093/nar/gkae410 %J Nucleic Acids Research.

29. Portero LR, Alonso-Reyes DG, Zannier F, Vazquez MP, Farías ME, Gärtner W, et al. Photolyases and Cryptochromes in UV-resistant Bacteria from High-altitude Andean Lakes. 2019;95(1):315–30. doi: 10.1111/php.13061.

30. Lamprecht-Grandío M, Cortesão M, Mirete S, de la Cámara MB, de Figueras CG, Pérez-Pantoja D, et al. Novel Genes Involved in Resistance to Both Ultraviolet Radiation and Perchlorate From the Metagenomes of Hypersaline Environments. Frontiers in microbiology. 2020;11:453. Epub 2020/04/16. doi: 10.3389/fmicb.2020.00453. PubMed PMID: 32292392; PubMed Central PMCID: PMCPMC7135895.

31. Maslowska KH, Makiela-Dzbenska K, Fijalkowska IJ. The SOS system: A complex and tightly regulated response to DNA damage. Environmental and molecular mutagenesis. 2019;60(4):368–84. Epub 2018/11/18. doi: 10.1002/em.22267. PubMed PMID: 30447030; PubMed Central PMCID: PMCPMC6590174.

32. Janion C. Inducible SOS response system of DNA repair and mutagenesis in Escherichia coli. International journal of biological sciences. 2008;4(6):338–44. Epub 2008/10/01. doi: 10.7150/ijbs.4.338. PubMed PMID: 18825275; PubMed Central PMCID: PMCPMC2556049.

33. Muntyan VS, Roumiantseva ML. Molecular Phylogenetic Analysis of Salt-Tolerance-Related Genes in Root-Nodule Bacteria Species Sinorhizobium meliloti. 2022;12(8):1968. PubMed PMID: doi:10.3390/agronomy12081968.

34. Godard T, Zühlke D, Richter G, Wall M, Rohde M, Riedel K, et al. Metabolic Rearrangements Causing Elevated Proline and Polyhydroxybutyrate Accumulation During the Osmotic Adaptation Response of Bacillus megaterium. Frontiers in bioengineering and biotechnology. 2020;8:47. Epub 2020/03/13. doi: 10.3389/fbioe.2020.00047. PubMed PMID: 32161752; PubMed Central PMCID: PMCPMC7053513.

35. Rath H, Reder A, Hoffmann T, Hammer E, Seubert A, Bremer E, et al. Management of Osmoprotectant Uptake Hierarchy in Bacillus subtilis via a SigB-Dependent Antisense RNA. Frontiers in microbiology. 2020;11:622. Epub 2020/05/07. doi: 10.3389/fmicb.2020.00622. PubMed PMID: 32373088; PubMed Central PMCID: PMCPMC7186363.

36. Blin K, Shaw S, Vader L, Szenei J, Reitz ZL, Augustijn HE, et al. antiSMASH 8.0: extended gene cluster detection capabilities and analyses of chemistry, enzymology, and regulation. Nucleic Acids Res. 2025;53(W1):W32–w8. Epub 2025/04/25. doi: 10.1093/nar/gkaf334. PubMed PMID: 40276974; PubMed Central PMCID: PMCPMC12230676 Italy and consults for Corteva Agriscience, Indianapolis, IN, USA. M.H.M. is a member of the Scientific Advisory Board of Hexagon Bio. All other authors declare to have no conflicts of interest.

37. Rebollar EA, Avitia M, Eguiarte LE, González-González A, Mora L, Bonilla-Rosso G, et al. Water–sediment niche differentiation in ancient marine lineages of Exiguobacterium endemic to the Cuatro Cienegas Basin. Environmental Microbiology. 2012;14(9):2323–33. doi: 10.1111/j.1462-2920.2012.02784.x.

38. Galván FS, Alonso-Reyes DG, Albarracín VH. From Genes to Nanotubes: Exploring the UV-Resistome in the Andean Extremophile *Exiguobacterium* Sp. S17. 2024:2024.06.28.600890. doi: 10.1101/2024.06.28.600890 %J bioRxiv.

39. Paredes Contreras BV, Vermelho AB, Casanova L, de Alencar Santos Lage C, Spindola Vilela CL, da Silva Cardoso V, et al. Enhanced UV-B photoprotection activity of carotenoids from the novel *Arthrobacter* sp. strain LAPM80 isolated from King George Island, Antarctica. Heliyon. 2025;11(1). doi: 10.1016/j.heliyon.2024.e41400.

40. Burbano-Erazo E, Ezquerro M, Sanchez-Bel P, Rodriguez-Concepcion M. Specific sets of geranylgeranyl diphosphate synthases and phytoene synthases control the production of carotenoids and ABA in different tomato tissues. Physiologia plantarum. 2025;177(1):e70052. Epub 2025/01/17. doi: 10.1111/ppl.70052. PubMed PMID: 39821357; PubMed Central PMCID: PMCPMC11738847.

41. Younes S, Al-Sulaiti A, Nasser EAA, Najjar H, Kamareddine L. Drosophila as a Model Organism in Host-Pathogen Interaction Studies. Frontiers in cellular and infection microbiology. 2020;10:214. Epub 2020/07/14. doi: 10.3389/fcimb.2020.00214. PubMed PMID: 32656090; PubMed Central PMCID: PMCPMC7324642.

42. Limmer S, Quintin J, Hetru C, Ferrandon D. Virulence on the fly: Drosophila melanogaster as a model genetic organism to decipher host-pathogen interactions. Current drug targets. 2011;12(7):978–99. Epub 2011/03/04. doi: 10.2174/138945011795677818. PubMed PMID: 21366519.

43. Coluzzi C, Garcillán-Barcia MP, de la Cruz F, Rocha EPC. Evolution of Plasmid Mobility: Origin and Fate of Conjugative and Nonconjugative Plasmids. Molecular Biology and Evolution. 2022;39(6). doi: 10.1093/molbev/msac115.

44. Garcillán-Barcia MP, Francia MV, de la Cruz F. The diversity of conjugative relaxases and its application in plasmid classification. FEMS microbiology reviews. 2009;33(3):657–87. Epub 2009/04/28. doi: 10.1111/j.1574-6976.2009.00168.x. PubMed PMID: 19396961.

45. Shirasu K, Kado CI. Membrane location of the Ti plasmid VirB proteins involved in the biosynthesis of a pilin-like conjugative structure on Agrobacterium tumefaciens. FEMS microbiology letters. 1993;111(2-3):287–94. Epub 1993/08/01. doi: 10.1111/j.1574-6968.1993.tb06400.x. PubMed PMID: 8405938.

46. Llosa M, Alkorta I. Coupling Proteins in Type IV Secretion. Current topics in microbiology and immunology. 2017;413:143–68. Epub 2017/01/01. doi: 10.1007/978-3-319-75241-9_6. PubMed PMID: 29536358.

47. Muschiol S, Erlendsson S, Aschtgen MS, Oliveira V, Schmieder P, de Lichtenberg C, et al. Structure of the competence pilus major pilin ComGC in Streptococcus pneumoniae. The Journal of biological chemistry. 2017;292(34):14134–46. Epub 2017/07/01. doi: 10.1074/jbc.M117.787671. PubMed PMID: 28659339; PubMed Central PMCID: PMCPMC5572924.

48. Damke PP, Celma L, Kondekar SM, Di Guilmi AM, Marsin S, Dépagne J, et al. ComFC mediates transport and handling of single-stranded DNA during natural transformation. Nature communications. 2022;13(1):1961. doi: 10.1038/s41467-022-29494-z.

49. Diallo A, Foster HR, Gromek KA, Perry TN, Dujeancourt A, Krasteva PV, et al. Bacterial transformation: ComFA is a DNA-dependent ATPase that forms complexes with ComFC and DprA. Molecular microbiology. 2017;105(5):741–54. Epub 2017/06/16. doi: 10.1111/mmi.13732. PubMed PMID: 28618091.

50. Takeno M, Taguchi H, Akamatsu T. Role of ComFA in controlling the DNA uptake rate during transformation of competent Bacillus subtilis. Journal of bioscience and bioengineering. 2011;111(6):618–23. Epub 2011/03/15. doi: 10.1016/j.jbiosc.2011.02.006. PubMed PMID: 21397556.

51. Kadlec K, Schwarz S. Novel ABC transporter gene, vga(C), located on a multiresistance plasmid from a porcine methicillin-resistant Staphylococcus aureus ST398 strain. Antimicrobial agents and chemotherapy. 2009;53(8):3589–91. Epub 2009/05/28. doi: 10.1128/aac.00570-09. PubMed PMID: 19470508; PubMed Central PMCID: PMCPMC2715595.

52. Peela SCM, Basu S, Sharma J, AlAsmari AF, AlAsmari F, Alalmaee S, et al. Structure Elucidation and Interaction Dynamics of MefA-MsrD Efflux Proteins in Streptococcus pneumoniae: Impact on Macrolide Susceptibility. ACS Omega. 2023;8(42):39454–67. doi: 10.1021/acsomega.3c05210.

53. Pawlowski AC, Stogios PJ, Koteva K, Skarina T, Evdokimova E, Savchenko A, et al. The evolution of substrate discrimination in macrolide antibiotic resistance enzymes. Nature communications. 2018;9(1):112. Epub 2018/01/11. doi: 10.1038/s41467-017-02680-0. PubMed PMID: 29317655; PubMed Central PMCID: PMCPMC5760710.

54. Pindi PK, Raghuveer Yadav P, Shiva Shanker A. Identification of Opportunistic Pathogenic Bacteria in Drinking Water Samples of Different Rural Health Centers and Their Clinical Impacts on Humans. 2013;2013(1):348250. doi: 10.1155/2013/348250.

55. Selvarajan R, Sibanda T, Pandian J, Mearns K. Taxonomic and Functional Distribution of Bacterial Communities in Domestic and Hospital Wastewater System: Implications for Public and Environmental Health. Antibiotics (Basel, Switzerland). 2021;10(9). Epub 2021/09/29. doi: 10.3390/antibiotics10091059. PubMed PMID: 34572642; PubMed Central PMCID: PMCPMC8470611.

56. Wang Q, Jiang G, Sun Z, Liang Y, Liu F, Shi J. Water quality and microecosystem of water tanks in karst mountainous area, Southwest China. Environmental Science and Pollution Research. 2024;31(9):12948–65. doi: 10.1007/s11356-024-31959-1.

57. Pavez VB, Pacheco N, Castro-Severyn J, Pardo-Esté C, Álvarez J, Zepeda P, et al. Characterization of biofilm formation by Exiguobacterium strains in response to arsenic exposure. Microbiology spectrum. 2023;11(6):e0265723. Epub 2023/10/11. doi: 10.1128/spectrum.02657-23. PubMed PMID: 37819075; PubMed Central PMCID: PMCPMC10714750.

58. Gao F, Liang Q, Zong R, Xie Y, Zhao C, Yang Y, et al. Isolation, Identification, and Antimicrobial Susceptibility of Exiguobacterium mexicanum from a Giraffe. Veterinary sciences. 2025;12(10). Epub 2025/10/28. doi: 10.3390/vetsci12100969. PubMed PMID: 41150109; PubMed Central PMCID: PMCPMC12568232.

