## SUPPLEMENTAL MATERIAL for "Environmental Selection Shapes Resistance, Metabolic, and Adaptive Capabilities in Exiguobacterium"

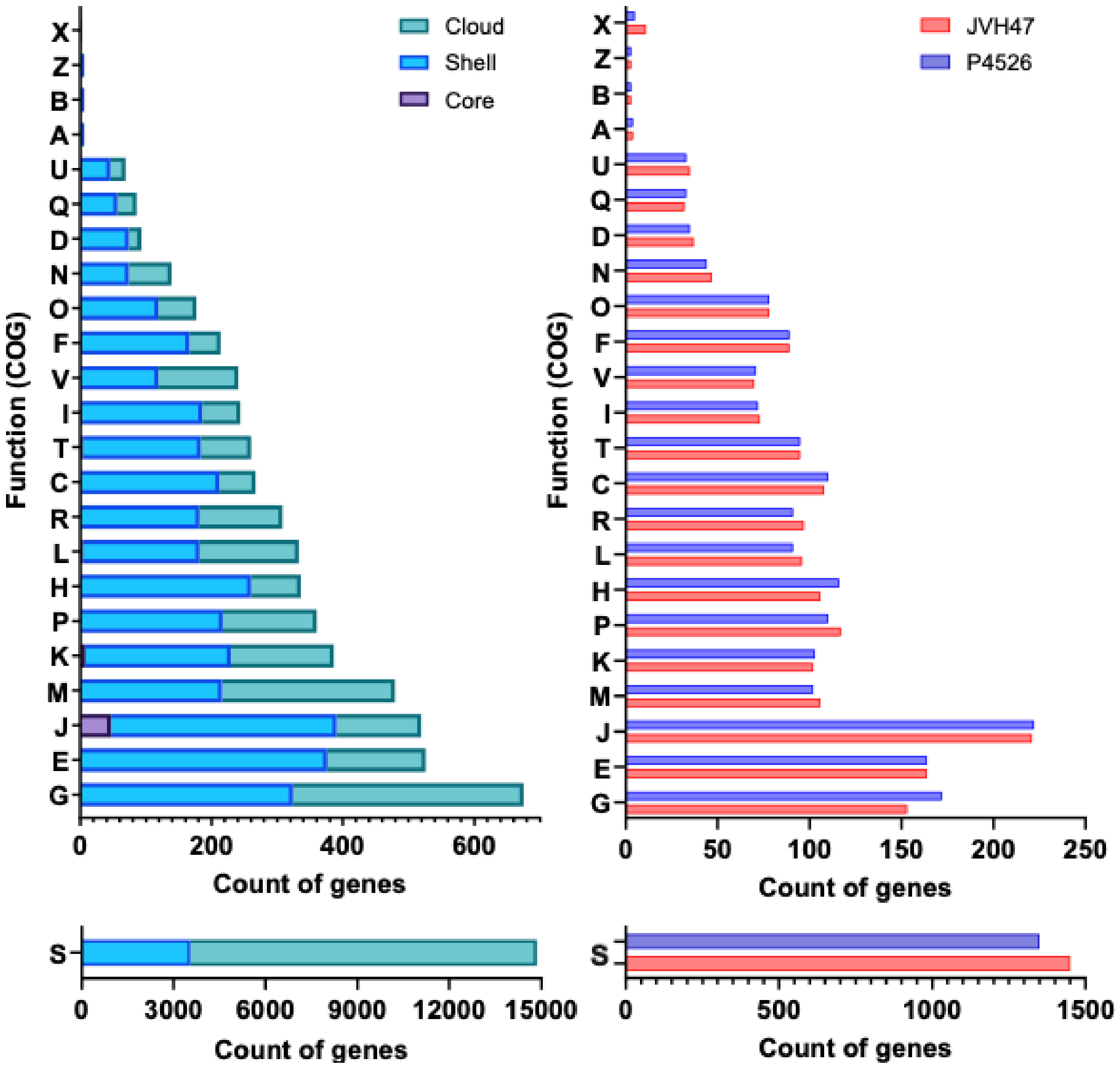

**Fig S1.** Function annotation of the ORFs present in *Exiguobacterium* strains according to Clusters of Orthologous Genes (COG). A gene could be assigned to more than one COG functional category. One letter COG code represent the following functions, A: RNA processing and modification, B: Chromatin structure and dynamics, C: Energy production and conversion, D: Cell cycle control, cell division, chromosome partitioning, E: Amino acid transport and metabolism, F: Nucleotide transport and metabolism, G: Carbohydrate transport and metabolism, H: Coenzyme transport and metabolism, I: Lipid transport and metabolism, J: Translation, ribosomal structure and biogenesis, K: Transcription, L: Replication, recombination and repair, M: Cell wall/membrane/envelope biogenesis, N: Cell motility, O: Posttranslational modification, protein turnover, chaperones, P: Inorganic ion transport and metabolism, Q: Secondary metabolites biosynthesis, transport and catabolism, R: General function prediction only, S: Function unknown, T: Signal transduction mechanisms, U: Intracellular trafficking, secretion, and vesicular transport, V: Defense mechanisms, X: Mobilome: prophages, transposons, Z: Cytoskeleton.

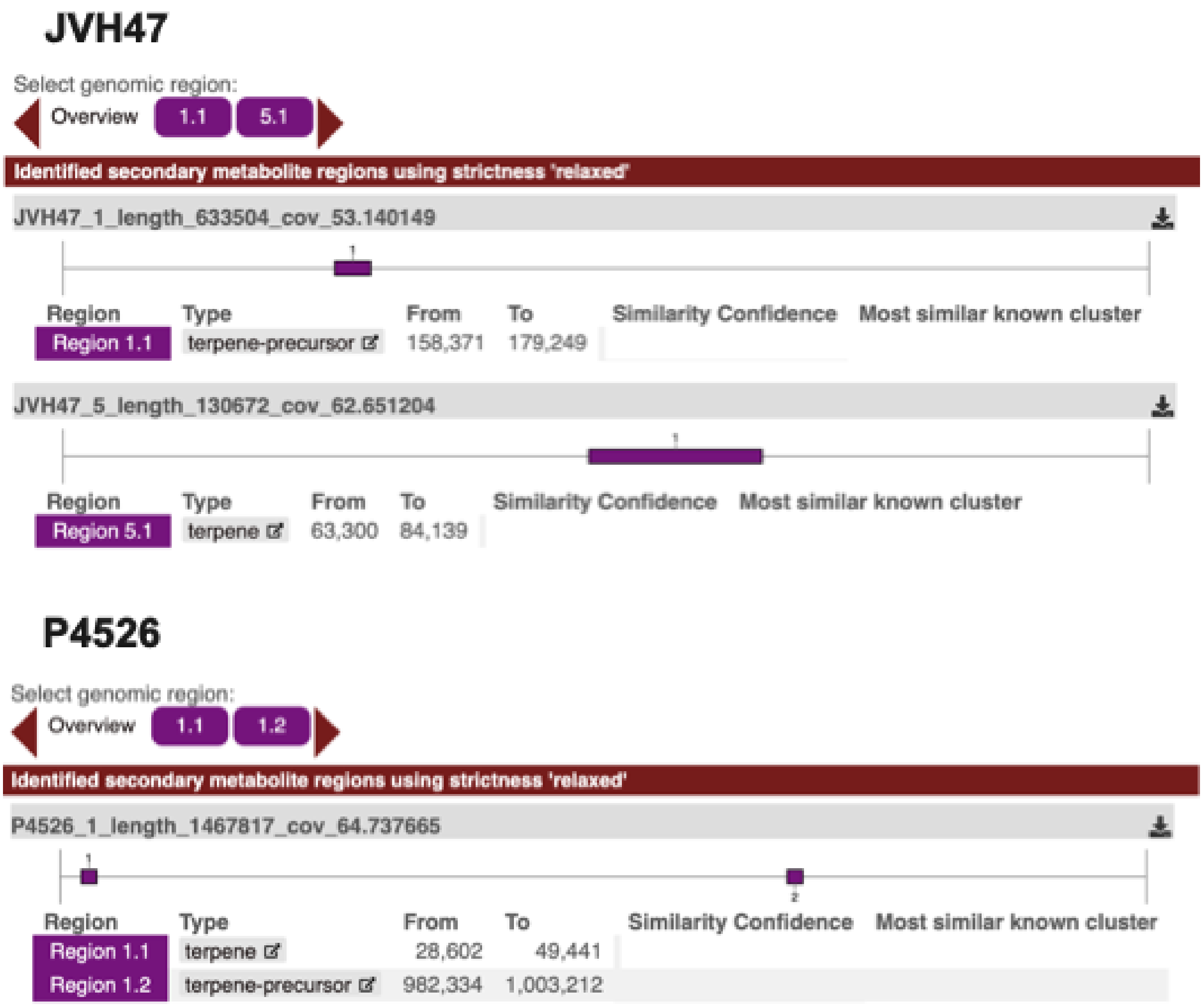

**Fig S2.** Prediction of biosynthetic gene clusters (BGC) *Exiguobacterium* strains JVH47 and P4526. Analysis was made using the online tool antiSMASH.

**Table S1.** Quality assessment of the assembly of the genomes of *Exiguobacterium* strains JVH47 and P4526.

| Statistics | JVH47 | P4526 |
|---|---|---|
| # contigs | 49 | 22 |
| # contigs (>= 0 bp) | 83 | 83 |
| # contigs (>= 1000 bp) | 46 | 18 |
| # contigs (>= 5000 bp) | 29 | 15 |
| # contigs (>= 10000 bp) | 26 | 13 |
| # contigs (>= 25000 bp) | 22 | 12 |
| # contigs (>= 50000 bp) | 18 | 10 |
| Largest contig | 633,504 | 1,467,817 |
| Total length | 2,970,744 | 2,871,014 |
| Total length (>= 0 bp) | 2,978,643 | 2,880,166 |
| Total length (>= 1000 bp) | 2,968,719 | 2,868,491 |
| Total length (>= 5000 bp) | 2,935,345 | 2,859,987 |
| Total length (>= 10000 bp) | 2,918,102 | 2,847,624 |
| Total length (>= 25000 bp) | 2,857,906 | 2,835,218 |
| Total length (>= 50000 bp) | 2,687,646 | 2,757,525 |
| GC (%) | 51.97 | 52.46 |
| N50 | 130,672 | 1,467,817 |
| N90 | 50,676 | 101,246 |
| auN | 265,010.8 | 835,236.6 |
| L50 | 5 | 1 |
| L90 | 18 | 8 |
| # N's per 100 kbp | 0 | 0 |

All statistics were calculated based on contigs of size ≥500 bp except when the size is specified

**Table S2.** Genes conferring resistance to extreme conditions and antibiotics in *Exiguobacterium* strains JVH47 and P4526. ORFs are annotated according to KEGG orthology.

| ***Exiguobacterium sp.* JVH47** | | |
|---|---|---|
| ***Osmosis resistance (non-unique gene: annotation)*** | ***UV resistance (non-unique gene: annotation)*** | ***AMR/mobile elements (non-unique gene: annotation)*** |
| opuD, betL: glycine betaine transporter | tag: DNA-3-methyladenine glycosylase I [EC:3.2.2.20] | comGC: competence protein ComGC |
| dps: starvation-inducible DNA-binding protein | sbcC, rad50: DNA repair protein SbcC/Rad50 | comGB: competence protein ComGB |
| sufB: Fe-S cluster assembly protein SufB | lexA: repressor LexA [EC:3.4.21.88] | comGA: competence protein ComGA |
| iscU, nifU: nitrogen fixation protein NifU and related proteins | recA: recombination protein RecA | comEC: competence protein ComEC |
| sufS: cysteine desulfurase / selenocysteine lyase [EC:2.8.1.7 4.4.1.16] | dinF, mepA, vmrA: MATE family, multidrug efflux pump | comEA: competence protein ComEA |
| sufD: Fe-S cluster assembly protein SufD | recN: DNA repair protein RecN (Recombination protein N) | comK: competence protein ComK |
| sufC: Fe-S cluster assembly ATP-binding protein | recO: DNA repair protein RecO (recombination protein O) | comFC: competence protein ComFC |
| PRODH, fadM, putB: proline dehydrogenase [EC:1.5.5.2] | mutM, fpg: formamidopyrimidine-DNA glycosylase [EC:3.2.2.23 4.2.99.18] | comFA: competence protein ComFA |
| galE, GALE: UDP-glucose 4-epimerase [EC:5.1.3.2] | mutT, NUDT15, MTH2: 8-oxo-dGTP diphosphatase [EC:3.6.1.55] | comC: leader peptidase (prepilin peptidase) / N-methyltransferase [EC:3.4.23.43 2.1.1.-] |
| kch, trkA, mthK, pch: voltage-gated potassium channel | uvrA: excinuclease ABC subunit A | comEB: dCMP deaminase [EC:3.5.4.12] |
| trpA: tryptophan synthase alpha chain [EC:4.2.1.20] | uvrB: excinuclease ABC subunit B | K07496: putative transposase |
| trpB: tryptophan synthase beta chain [EC:4.2.1.20] | alkA: DNA-3-methyladenine glycosylase II [EC:3.2.2.21] | K07496: putative transposase |
| trpC: indole-3-glycerol phosphate synthase [EC:4.1.1.48] | ogt, MGMT: methylated-DNA-[protein]-cysteine S-methyltransferase [EC:2.1.1.63] | K07496: putative transposase |
| trkH, trkG, ktrB, ktrD: trk/ktr system potassium uptake protein | ahpC: NADH-dependent peroxiredoxin subunit C [EC:1.11.1.26] | K07496: putative transposase |
| ftsZ: cell division protein FtsZ | mutT, NUDT15, MTH2: 8-oxo-dGTP diphosphatase [EC:3.6.1.55] | K07496: putative transposase |
| csoR, ricR: CsoR family transcriptional regulator, | mutS2: DNA mismatch repair protein MutS2 | K07497: putative transposase |

| | | |
|---|---|---|
| copper-sensing transcriptional repressor | | |
| copA, ctpA, ATP7: P-type Cu+ transporter [EC:7.2.2.8] | UNG, UDG: uracil-DNA glycosylase [EC:3.2.2.27] | K07482: transposase, IS30 family |
| ATOX1, ATX1, copZ, golB: copper chaperone | recQ: ATP-dependent DNA helicase RecQ [EC:5.6.2.4] | |
| | | K07497: putative transposase |
| | ftsK, spoIIIE: DNA segregation ATPase FtsK/SpoIIIE, S-DNA-T family | |
| fixA, etfB: electron transfer flavoprotein beta subunit | | |
| | | K07497: putative transposase |
| fixB, etfA: electron transfer flavoprotein alpha subunit | recG: ATP-dependent DNA helicase RecG [EC:5.6.2.4] | |
| | | K07497: putative transposase |
| | ogt, MGMT: methylated-DNA-[protein]-cysteine S-methyltransferase [EC:2.1.1.63] | |
| | | blaTEM-116: class A broad-spectrum beta-lactamase TEM-116 |
| trkA, ktrA, ktrC: trk/ktr system potassium uptake protein | | |
| perR: Fur family transcriptional regulator, peroxide stress response regulator | | |
| | dinB: DNA polymerase IV [EC:2.7.7.7] | mphN: macrolide 2'-phosphotransferase MphN |
| trkH, trkG, ktrB, ktrD: trk/ktr system potassium uptake protein | | |
| | dinF, mepA, vmrA: MATE family, multidrug efflux pump | |
| | | MOBP2: |
| patB, malY: cysteine-S-conjugate beta-lyase [EC:4.4.1.13] | | |
| | mutS: DNA mismatch repair protein MutS | |
| ATOX1, ATX1, copZ, golB: copper chaperone | mutL: DNA mismatch repair protein MutL | |
| ATOX1, ATX1, copZ, golB: copper chaperone | osmC, ohr: lipoyl-dependent peroxiredoxin [EC:1.11.1.28] | |
| copA, ctpA, ATP7: P-type Cu+ transporter [EC:7.2.2.8] | mutS2: DNA mismatch repair protein MutS2 | |
| ATOX1, ATX1, copZ, golB: copper chaperone | uvrC: excinuclease ABC subunit C | |
| yclQ, ceuA: iron-siderophore transport system substrate-binding protein | | |
| | pilT: twitching motility protein PilT | |
| yclN, ceuB: iron-siderophore transport system permease protein | | |
| | radC: DNA repair protein RadC | |
| yclO, ceuC: iron-siderophore transport system permease protein | mutT, NUDT15, MTH2: 8-oxo-dGTP diphosphatase [EC:3.6.1.55] | |
| yclP, ceuD: iron-siderophore transport system ATP-binding protein [EC:7.2.2.-] | | |
| | ruvA: holliday junction DNA helicase RuvA | |
| panD: aspartate 1-decarboxylase [EC:4.1.1.11] | ruvB: holliday junction DNA helicase RuvB [EC:5.6.2.4] | |
| | recJ: single-stranded-DNA-specific exonuclease [EC:3.1.-.-] | |
| panC: pantoate--beta-alanine ligase [EC:6.3.2.1] | | |

| | |
|---|---|
| panB: 3-methyl-2-oxobutanoate hydroxymethyltransferase [EC:2.1.2.11]<br>galE, GALE: UDP-glucose 4-epimerase [EC:5.1.3.2]<br><br>rex: redox-sensing transcriptional repressor<br><br>ATOX1, ATX1, copZ, golB: copper chaperone | recX: regulatory protein<br>mutY: A/G-specific adenine glycosylase [EC:3.2.2.31]<br>perR: Fur family transcriptional regulator, peroxide stress response regulator<br>uvrD, pcrA: ATP-dependent DNA helicase UvrD/PcrA [EC:5.6.2.4]<br>mutT, NUDT15, MTH2: 8-oxo-dGTP diphosphatase [EC:3.6.1.55]<br>mutT, NUDT15, MTH2: 8-oxo-dGTP diphosphatase [EC:3.6.1.55]<br>radC: DNA repair protein RadC<br>uvrD, pcrA: ATP-dependent DNA helicase UvrD/PcrA [EC:5.6.2.4]<br>ssb: single-strand DNA-binding protein<br>recF: DNA replication and repair protein RecF<br>mutT, NUDT15, MTH2: 8-oxo-dGTP diphosphatase [EC:3.6.1.55]<br>radA, sms: DNA repair protein RadA/Sms<br>recU: recombination protein U<br>NTHL1, nth: endonuclease III [EC:3.2.2.- 4.2.99.18]<br>dinG: ATP-dependent DNA helicase DinG [EC:5.6.2.3]<br>mfd: transcription-repair couplingfactor (superfamily II helicase) [EC:5.6.2.4]<br>radC: DNA repair protein RadC<br>tadA: tRNA(adenine34) deaminase [EC:3.5.4.33]<br>recR: recombination protein RecR |

### *Exiguobacterium sp.* P4526

| ***Osmosis resistance (non-unique gene: annotation)*** | ***UV resistance (non-unique gene: annotation)*** | ***AMR/mobile elements (non-unique gene: annotation)*** |
|---|---|---|

| Non-unique Gene name: Annotation | Non-unique Gene name: Annotation | Non-unique Gene name: Annotation |
|---|---|---|
| csoR, ricR: CsoR family transcriptional regulator, copper-sensing transcriptional repressor<br>copA, ctpA, ATP7: P-type Cu+ transporter [EC:7.2.2.8]<br>ATOX1, ATX1, copZ, golB: copper chaperone<br>fixA, etfB: electron transfer flavoprotein beta subunit<br>fixB, etfA: electron transfer flavoprotein alpha subunit<br>trkA, ktrA, ktrC: trk/ktr system potassium uptake protein<br>ftsZ: cell division protein FtsZ<br>panB: 3-methyl-2-oxobutanoate hydroxymethyltransferase [EC:2.1.2.11]<br>panC: pantoate--beta-alanine ligase [EC:6.3.2.1]<br>panD: aspartate 1-decarboxylase [EC:4.1.1.11]<br>patB, malY: cysteine-S-conjugate beta-lyase [EC:4.4.1.13]<br>opuD, betL: glycine betaine transporter<br>dps: starvation-inducible DNA-binding protein<br>sufB: Fe-S cluster assembly protein SufB<br>iscU, nifU: nitrogen fixation protein NifU and related proteins<br>sufS: cysteine desulfurase / selenocysteine lyase [EC:2.8.1.7 4.4.1.16]<br>sufD: Fe-S cluster assembly protein SufD<br>sufC: Fe-S cluster assembly ATP-binding protein<br>PRODH, fadM, putB: proline dehydrogenase [EC:1.5.5.2]<br>opuD, betL: glycine betaine transporter | dinF, mepA, vmrA: MATE family, multidrug efflux pump<br>osmC, ohr: lipoyl-dependent peroxiredoxin [EC:1.11.1.28]<br>mutS2: DNA mismatch repair protein MutS2<br>uvrC: excinuclease ABC subunit C<br>pilT: twitching motility protein PilT<br>radC: DNA repair protein RadC<br>mutT, NUDT15, MTH2: 8-oxo-dGTP diphosphatase [EC:3.6.1.55]<br>ruvA: holliday junction DNA helicase RuvA<br>ruvB: holliday junction DNA helicase RuvB [EC:5.6.2.4]<br>recJ: single-stranded-DNA-specific exonuclease [EC:3.1.-.-]<br>recG: ATP-dependent DNA helicase RecG [EC:5.6.2.4]<br>ftsK, spoIIIE: DNA segregation ATPase FtsK/SpoIIIE, S-DNA-T family<br>recQ: ATP-dependent DNA helicase RecQ [EC:5.6.2.4]<br>dinG: ATP-dependent DNA helicase DinG [EC:5.6.2.3]<br>NTHL1, nth: endonuclease III [EC:3.2.2.- 4.2.99.18]<br>recU: recombination protein U<br>mutS: DNA mismatch repair protein MutS<br>mutL: DNA mismatch repair protein MutL<br>mutT, NUDT15, MTH2: 8-oxo-dGTP diphosphatase [EC:3.6.1.55]<br>tag: DNA-3-methyladenine glycosylase I [EC:3.2.2.20] | comC: leader peptidase (prepilin peptidase) / N-methyltransferase [EC:3.4.23.43 2.1.1.-]<br>comEB: dCMP deaminase [EC:3.5.4.12]<br>comGB: competence protein ComGB<br>comEC: competence protein ComEC<br>comEA: competence protein ComEA<br>comK: competence protein ComK<br>comFC: competence protein ComFC<br>comFA: competence protein ComFA<br>K07496: putative transposase<br>K07496: putative transposase<br>rayT: REP-associated tyrosine transposase<br>K07496: putative transposase<br>K07496: putative transposase |

| | |
|---|---|
| trpC: indole-3-glycerol phosphate synthase [EC:4.1.1.48]<br>trpB: tryptophan synthase beta chain [EC:4.2.1.20]<br>trpA: tryptophan synthase alpha chain [EC:4.2.1.20]<br>kch, trkA, mthK, pch: voltage-gated potassium channel<br><br>galE, GALE: UDP-glucose 4-epimerase [EC:5.1.3.2]<br><br>galE, GALE: UDP-glucose 4-epimerase [EC:5.1.3.2]<br><br>yclQ, ceuA: iron-siderophore transport system substrate-binding protein<br>yclN, ceuB: iron-siderophore transport system permease protein<br>yclO, ceuC: iron-siderophore transport system permease protein<br>yclP, ceuD: iron-siderophore transport system ATP-binding protein [EC:7.2.2.-]<br>rex: redox-sensing transcriptional repressor<br>perR: Fur family transcriptional regulator, peroxide stress response regulator<br>trkH, trkG, ktrB, ktrD: trk/ktr system potassium uptake protein<br>trkH, trkG, ktrB, ktrD: trk/ktr system potassium uptake protein | sbcC, rad50: DNA repair protein SbcC/Rad50<br>lexA: repressor LexA [EC:3.4.21.88]<br>recA: recombination protein RecA<br>dinF, mepA, vmrA: MATE family, multidrug efflux pump<br>recN: DNA repair protein RecN (Recombination protein N)<br>recO: DNA repair protein RecO (recombination protein O)<br>mutM, fpg: formamidopyrimidine-DNA glycosylase [EC:3.2.2.23 4.2.99.18]<br>mutT, NUDT15, MTH2: 8-oxo-dGTP diphosphatase [EC:3.6.1.55]<br><br>uvrA: excinuclease ABC subunit A<br><br>uvrB: excinuclease ABC subunit B<br>mutS2: DNA mismatch repair protein MutS2<br>mutT, NUDT15, MTH2: 8-oxo-dGTP diphosphatase [EC:3.6.1.55]<br>uvrD, pcrA: ATP-dependent DNA helicase UvrD/PcrA [EC:5.6.2.4]<br>ahpC: NADH-dependent peroxiredoxin subunit C [EC:1.11.1.26]<br>mutT, NUDT15, MTH2: 8-oxo-dGTP diphosphatase [EC:3.6.1.55]<br>ogt, MGMT: methylated-DNA-[protein]-cysteine S-methyltransferase [EC:2.1.1.63]<br>ssb: single-strand DNA-binding protein<br>recF: DNA replication and repair protein RecF<br>recX: regulatory protein |

| | |
|---|---|
| | mutY: A/G-specific adenine glycosylase [EC:3.2.2.31]<br>perR: Fur family transcriptional regulator, peroxide stress response regulator<br>UNG, UDG: uracil-DNA glycosylase [EC:3.2.2.27]<br>uvrD, pcrA: ATP-dependent DNA helicase UvrD/PcrA [EC:5.6.2.4]<br>radA, sms: DNA repair protein RadA/Sms<br>mfd: transcription-repair couplingfactor (superfamily II helicase) [EC:5.6.2.4]<br>dinB: DNA polymerase IV [EC:2.7.7.7]<br>tadA: tRNA(adenine34) deaminase [EC:3.5.4.33]<br>recR: recombination protein RecR |
